# Cas12a-based on-site and rapid nucleic acid detection of African swine fever

**DOI:** 10.1101/729590

**Authors:** Jing Bai, Haosi Lin, Haojian Li, Yang Zhou, Junshan Liu, Guorui Zhong, Luting Wu, Weifan Jiang, Hongli Du, Jinyi Yang, Qingmei Xie, Lizhen Huang

## Abstract

The mortality rate of hemorrhagic African swine fever (ASF), which targets domestic pigs and is caused by African swine fever virus (ASFV), can reach 100%. ASF has been reported in 25 Chinese provinces since August 2018. There is no effective treatment or vaccine for it and the present molecular diagnosis technologies have trade-offs in sensitivity, specificity, cost and speed, and none of them cater perfectly to ASF control. Thus, a technology that overcomes the need for laboratory facilities, is relatively low cost, and rapidly and sensitively detects ASFV would be highly valuable. Here, we describe an RAA-Cas12a-based system that combines recombinase-aided amplification (RAA) and CRISPR/Cas12a for ASFV detection. The fluorescence intensity readout of this system detected ASFV p72 gene levels as low as 10 aM. For on-site ASFV detection, lateral-flow strip readout was introduced for the first time in the RAA-Cas12a based system (named CORDS, **C**as12a-based **O**n-site and **R**apid **D**etection **S**ystem). We used CORDS to detect target DNA highly specifically using the lateral-flow strip readout. CORDS could identify the p72 gene at femtomolar sensitivity in an hour at 37°C, and only requires an incubator. For ease of use, the regents of CORDS was lyophilized to three tubes and remained the same sensitivity when stored at 4 °C for at least 7 days. Thus, CORDS provides a rapid, sensitive and easily operable method for ASFV on-site detection. Lyophilized CORDS can withstand long-term transportation and storage, and is ready for field applications.

## Introduction

African swine fever (ASF) is a highly contagious hemorrhagic disease of domestic pigs and wild boars. ASF is caused by a large, complex double-stranded DNA virus, the African swine fever virus (ASFV) ^[1]^. The disease shares similar clinical signs with classical swine fever (CSF) and several other swine diseases^[2]^, making its diagnosis difficult, especially outside of a diagnostics laboratory. Since the first ASF case emerged in China in August 2018, ASF has spread to 25 provinces and 114 ASF infection cases have been reported^[3]^. By January 2019 in China, 10,398 of 268,698 pigs infected with ASFV had died and the remainder were culled^[3]^. There is no effective treatment or vaccine against ASF yet, so controlling ASF mainly relies on animal slaughter and sanitary measures^[1, 4]^. Therefore, accurate and timely diagnosis of ASF infections is crucial for controlling epidemics.

The recommended gold standard for diagnosis according to the World Organization for Animal Health (OIE) is virus isolation. But this is time-consuming and labor-intensive, and virus isolation is not a method applicable to disease monitoring and control^[2]^. The antigen enzyme-linked immunosorbent assay is a rapid and convenient method for detecting ASFV antigens^[2]^. However, it lacks sensitivity in subacute cases or for early-stage infections and has poor specificity because it cannot correctly identify different viral strains^[2, 5]^. Hence, the molecular tools for detecting ASFV, such as polymerase chain reaction (PCR) and reverse-transcriptase (RT)-PCR, have become popular because of their sensitivity and specificity for ASF diagnosis^[2]^. Nevertheless, such methods are not suitable for on-site situations (e.g., on farms) or for rapid viral detection as they require thermal cycling instruments and skilled operators.^[5, 6]^. Isothermal amplification techniques, such as recombinase polymerase amplification (RPA), loop mediated isothermal amplification and cross-priming amplification, have been successfully used to detect ASFV^[2]^. Recently, many novel isothermal amplification assays in combination with immunochromatographic strips have also been developed for on-site detection of ASFV^[7, 8]^. With no need for instruments and skilled operators, isothermal amplification has become a promising on-site detection method. But like other amplification technologies, its resolution depends on the binding specificity between the primers and templates, which can limit their accuracy^[5]^.

With its ability to discriminate a single base mutation, the CRISPR/Cas system holds promise as a detection method^[9, 10]^. The trans-cleavage activity of the recently discovered Cas12a, Cas13a, and Cas14 systems in particular, helps them first bind to the target specifically and then amplify the binding signal by cutting the dual-labeled probe in a nonspecific way^[9-16]^. New gene detection technologies using the CRISPR/Cas12a and Cas13a systems, such as DETECTR, SHERLOCK and HOLMES, have extremely high sensitivities and specificities (single base resolution), and have been developed in combination with other gene amplification methods, including PCR and RPA^[11-15]^. Cas12-based DECTECTR and HOLMES need fluorescent detection equipment and therefore are not suitable for field tests^[14, 15]^. In contrast, when Cas13-based SHERLOCK is combined with an immunochromatographic strip, it becomes a good choice for on-site detection^[12, 13]^. However with SHERLOCK, both the target and probe are RNA, which increases the detection costs and reduces the assay’s stability. Moreover, the RNA probe can generate false positive results because RNase is ever present and RNA is generally unstable^9, 11-13]^. The newly reported naked-eye gene detection platform based on the CRISPR/Cas12a/13a system also needs a centrifugal machine and whole blood samples can interrupt the naked-eye detection^[17]^.

Therefore, the Cas12a-based system using a single-stranded DNA probe is more suitable for viral detection where limited conditions for on-site detection exist such as on farms^[9]^. In this study, we combined Cas12a with recombinase aided amplification (RAA) and an immunochromatographic lateral flow strip to develop a test strip method, termed CORDS (**C**as12a-based **O**n-site and **R**apid **D**etection **S**ystem) for ASFV on-site detection. CORDS can detect the ASFV target at a sensitivity of 1 fM within 1 h of initiation and without cross reaction to 13 porcine virus DNAs. When the RAA-Cas12a-based system was combined with fluorescence, the sensitivity increased to 10 aM. Finally, CORDS can also be freeze-dried in three tubes, making it suitable for transportation and storage, factors that are helpful in resource limited on-site situations. The lyophilized CORDS products are highly stable. Successful detection of ASFV with sensitivity comparable to that of SHERLOCK and without the need for expensive instruments makes our method a robust on-site detection platform.

## Results

### Cas12a cleavage activity guided by crRNA

Four targeted sequences were identified by NCBI BLAST and Perl language programming based on the TTTN-N18 rule for LbCas12a targeting, and there was a 20-bp (minimum) conserved sequence flanking both sides of the target within 250 bp. The target sequences are listed in Table 1. Based on these sequences, crRNAs were designed and prepared via *in vitro* transcription (Table 1). Target 1, which is located in the p72 gene, was used for subsequent tests. The *in vitro* cleavage results confirmed the crRNA guided LbCas12a cleavage activity of the assay (Fig. 1).

**Fig. 1.**
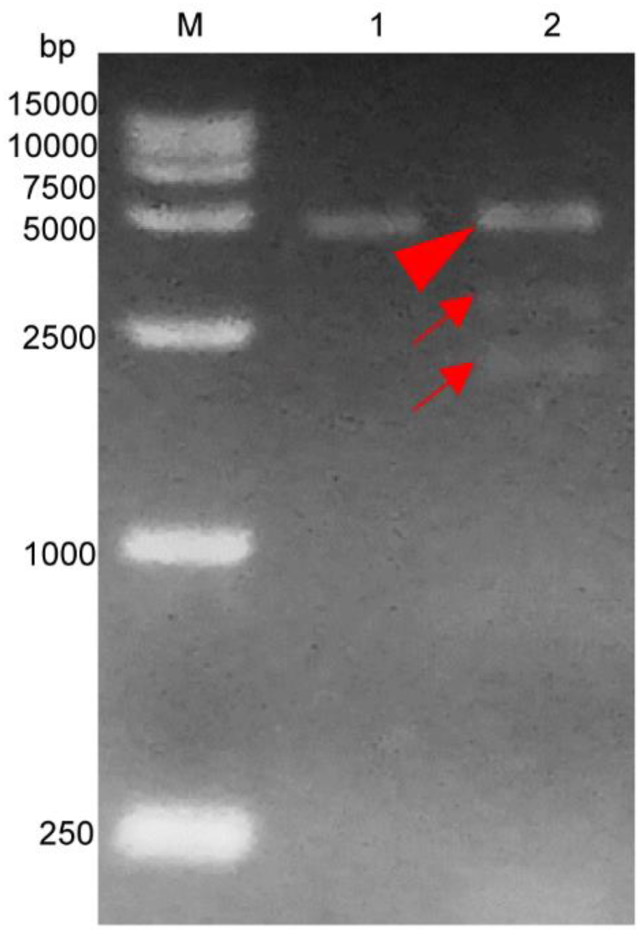
Cleavage activity of crRNA-guided LbCas12a. M: DL15000 DNA marker; 1: intact linearized dsDNA (4618 bp); 2: cleavage products from linearized dsDNA. Double-strand DNA (dsDNA) binding to the LbCas12a-crRNA complex is marked by a red triangle where the band can be seen to shift upwards compared with the intact linearized dsDNA. The expected cleavage products are marked by red arrows.

### The Cas12a-based ASFV nucleic acid fluorescence reporting system

We then established a fluorescence reporting system based on the cleavage activity of LbCas12a. To evaluate its validity, a negative control and pUC57–p72 were separately added to the system as targets. The statistical analysis revealed that there was no significant difference (p > 0.05) between the blank control and the negative control groups. The pUC57–p72 group showed significantly higher fluorescence intensity than the other groups (P < 0.0001). These results confirm that crRNA-complementary dsDNA was able to trigger LbCas12a to cleave the ssDNA FQ–labeled reporter (Fig. 2A). The reporting system had sufficient sensitivity to detect pUC57–p72 at a concentration of 1 × 10^-9^ M (Fig. 2B).

**Fig. 2.**
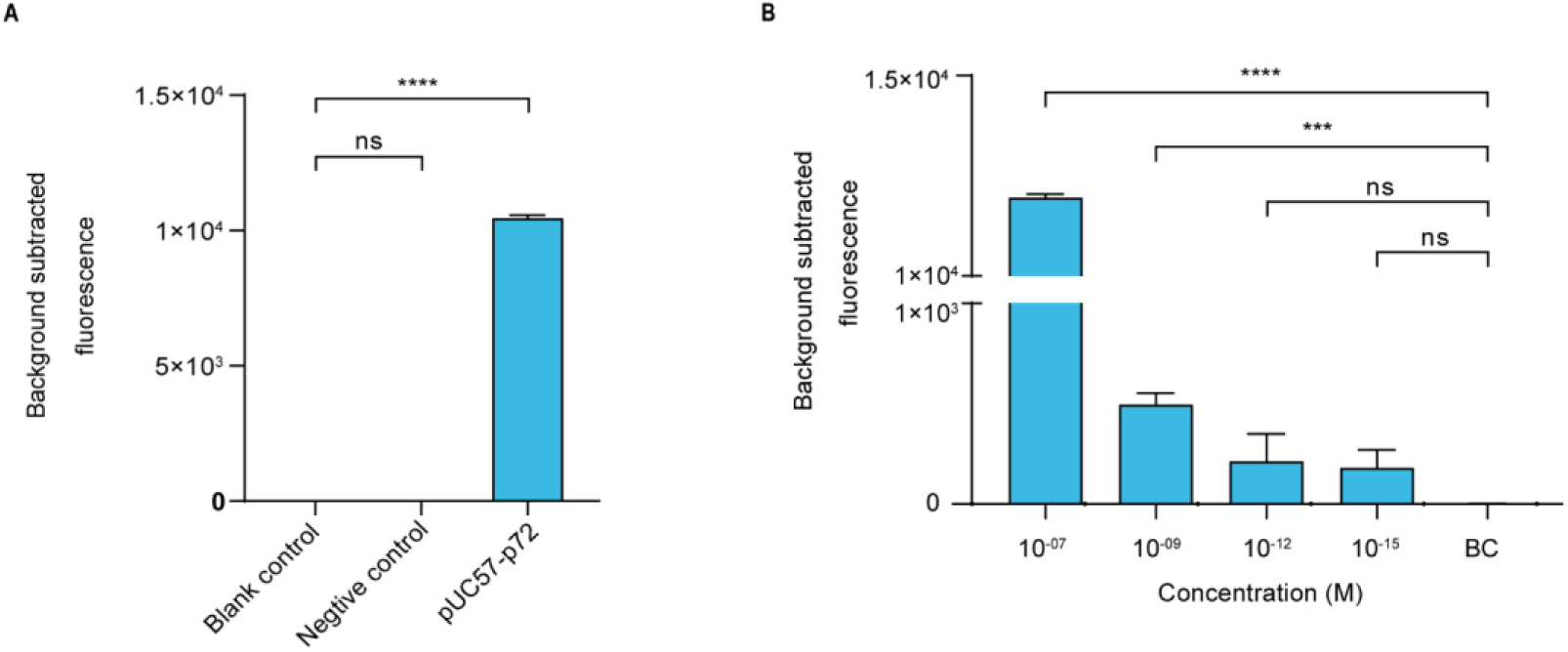
Cas12a-based ASFV NA fluorescence reporting system. **(A)**. Validation of the Cas12a-based ASFV nucleic acid fluorescence reporting system. The concentrations of the negative control and pUC57–p72 were both 1 × 10^-8^ M. **(B)**. Sensitivity of the Cas12a-based ASFV NA fluorescence reporting system. Background subtracted fluorescence was the fluorescence intensity of the experimental group against the blank control. Error bars in **(A)** and **(B)** represent the mean ± SD, where n = 3 replicates. ***p ≤ 0.001 and ****p ≤ 0.0001.

### Establishing the RAA-Cas12a-fluorescence assay

Before assay establishment, we optimized the concentration of LbCas12a by measuring the fluorescence intensity kinetics of LbCas12a targeting. The 50 nM group showed a more obvious difference in fluorescence intensity than the 250 nM group (Fig. 3A). Thus, the concentration of LbCas12a was set at 50 nM for the fluorescent assays in subsequent experiments. We also investigated the appropriate concentration ratio of LbCas12a to crRNA. The group with a 1:2 ratio showed better performance and reproducibility than the others (Fig. 3B). The optimized system was combined with RAA to establish the RAA-Cas12a-fluorescence assay. The LOD for the assay reached 10 aM (Fig. 3C).

**Fig. 3.**
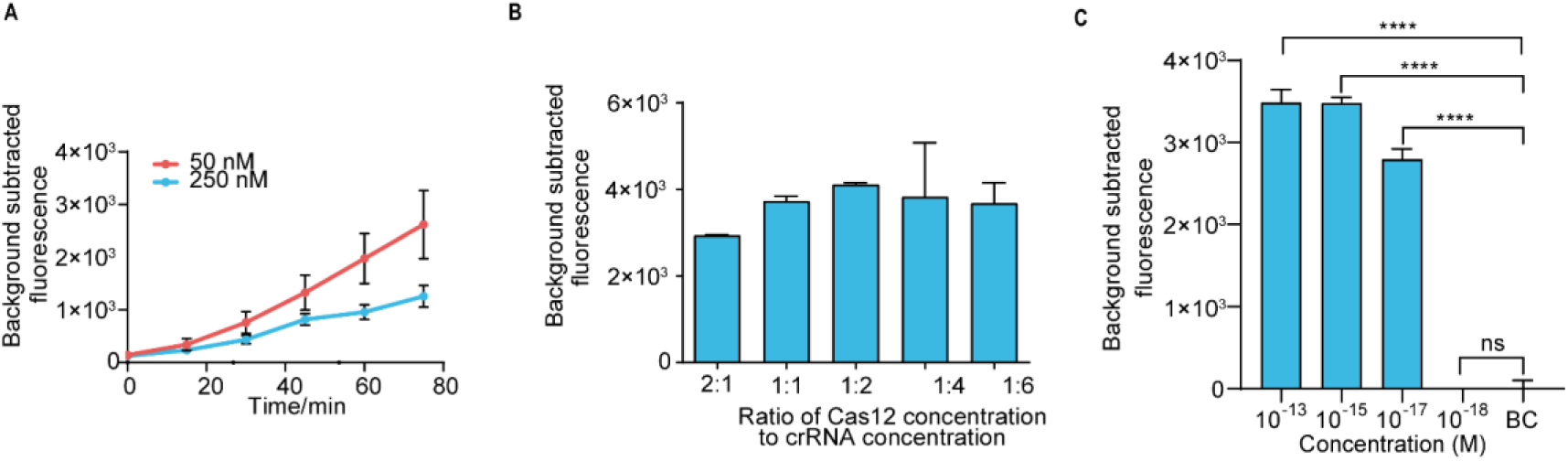
Establishing the RAA-Cas12a-fluorescence assay. **(A)**. Fluorescence kinetics at different LbCas12a concentrations. **(B)**. Optimizing the concentration ratio of LbCas12a to crRNA. **(C)**. Sensitivity of the RAA-Cas12a-fluorescence assay. Background subtracted fluorescence was the fluorescence intensity of the experimental group against the blank control. Error bars in **(A), (B)** and **(C)** represent the mean ± SD, where n = 3 replicates. *p ≤ 0.05 and ****p ≤ 0.0001.

### Establishing the CORDS assay

For on-site detection, we substituted the fluorescence intensity readout in the RAA-Cas12a-fluorescence assay with lateral flow strips to establish the CORDS assay (Fig. 4A). To determine the appropriate concentration of LbCas12a for the CORDS assay, we next compared two groups in which the concentrations of LbCas12a were 250 nM and 50 nM. The former showed a more obvious test-band intensity difference between the experiment group and the blank control (Fig. 4B). Therefore, we chose 250 nM as the concentration of LbCas12a for the CORDS assays, and the dsDNA targets were diluted from 1 × 10^-9^ M to 1 × 10^-17^ M for sensitivity evaluation. The LOD of the assay was 1 × 10^-15^ M according to the concentration gradient test results (Fig. 4C). The gray values of the test bands in the CORDS assays were measured and recorded. A positive result was recorded when the concentration of the dsDNA target was 1 × 10^-15^ M, which further confirmed the assay’s sensitivity (Fig. 4D). The 13 NA samples from typical porcine viruses were tested with this assay, all of which were negative. Thus, no cross-reactions were observed (Fig. 4E)

**Fig. 4.**
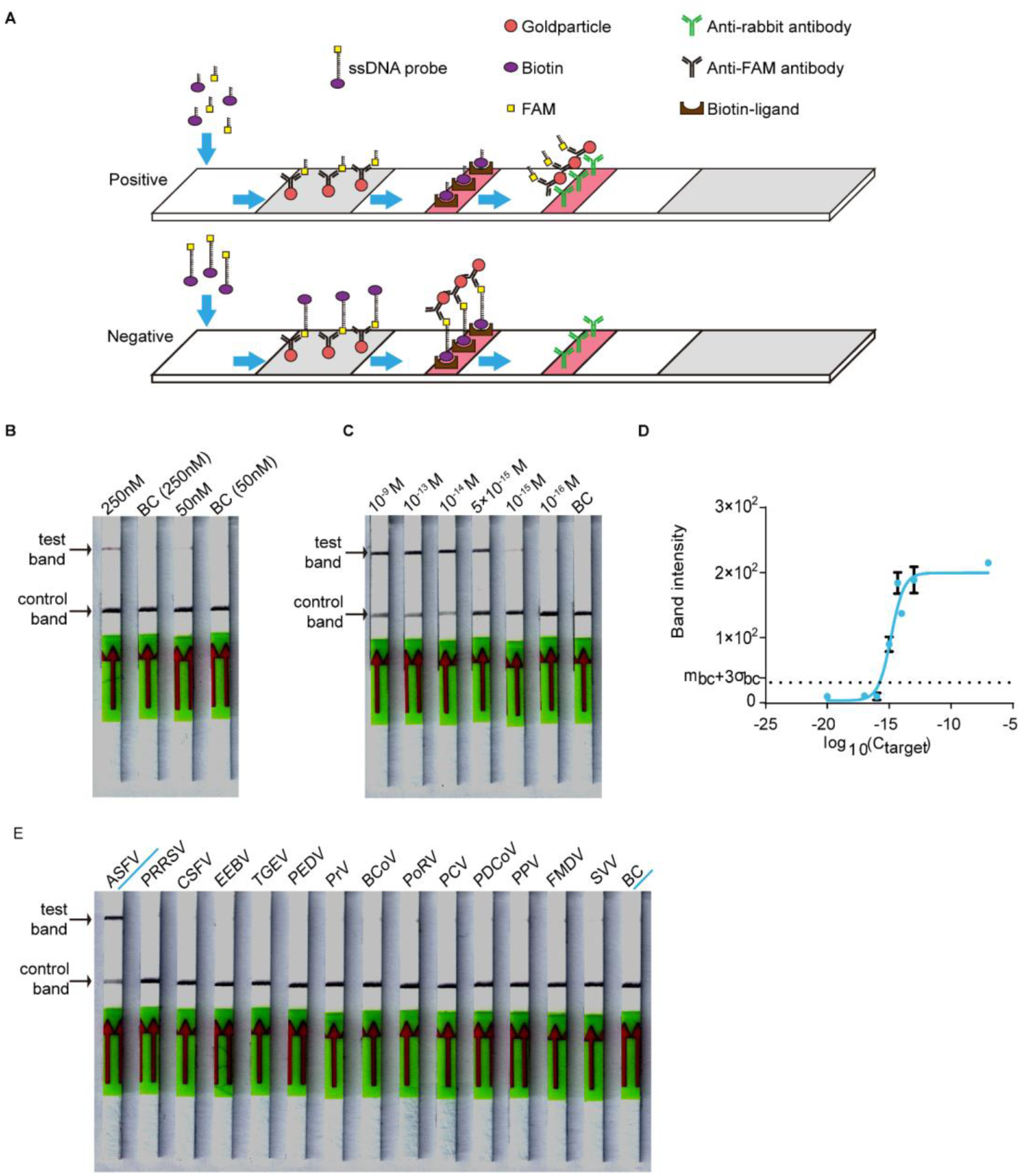
Establishing the CORDS assay. **(A)**. Schematic diagram of the CORDS assay. **(B)**. Optimization of the concentration of LbCas12a. **(C)**. Sensitivity of the CORDS assay. **(D)**. Mean gray values of the test band at different dsDNA target concentrations. The dashed line m_bc_+ 3σ_bc_ indicates the positive cut-off. **(E)**. Specificity of the CORDS assay. BC: blank control.

### Assessing the stability of the lyophilized CORDS assay

The components of the CORDS assay were lyophilized in three tubes to simplify the workflow (Fig. 5A). The lyophilized RAA buffer mix (tube B) and Cas mix (tube C) were verified and the two tubes were tested for further confirmation. The results illustrated that the LOD was still 1 × 10^-15^ M (Fig. 5B). Accelerated stability tests were performed on tubes B and C. Tube B was still functional at a target concentration of 1 × 10^-15^ M after storage at 37°C for 168 h (Fig. 5C). Tube C still worked well after storage at 4°C for 168 h without any sensitivity loss (Fig. 5D).

**Fig. 5.**
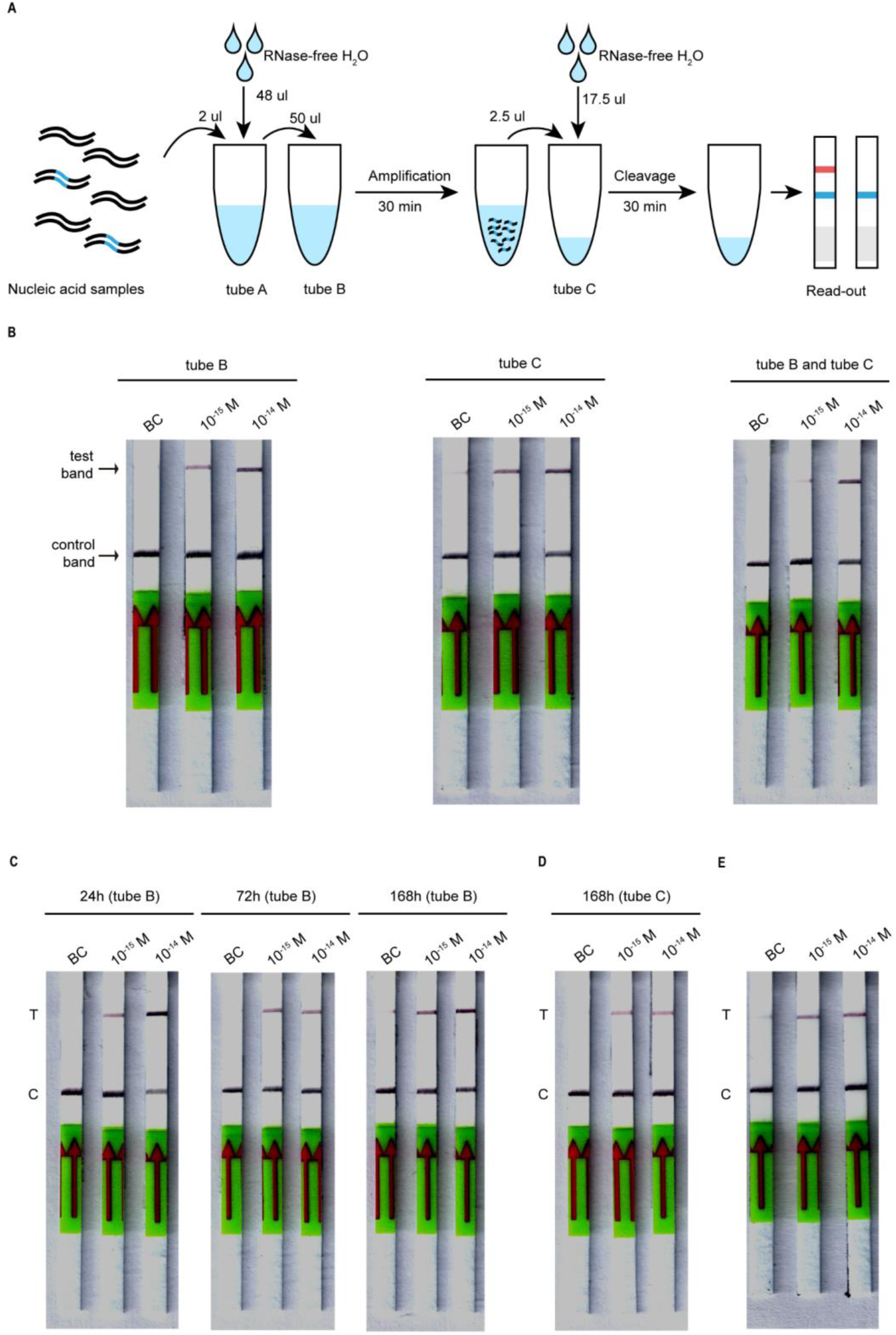
Lyophilized CORDS assay. **(A).** Workflow for the lyophilized CORDS assay. **(B).** Verification of tubes B and C. **(C).** Accelerated stability test on tube B at 37°C. **(D).** Accelerated stability test on tube C at 4°C. **(E).** Reduction of the incubation time for LbCas12a cleavage. T and C in (C), (D) and (E) represent the test and control bands, respectively.

We also adjusted the incubation time for LbCas12a cleavage, reducing it from 1 h to 30 min, and the detection time reduced correspondingly from 1.5 h to 1 h. The LOD still reached 1 × 10^-15^ M (Fig. 5E).

## Discussion

Since August 2018, ASF has affected 25 Chinese provinces and caused great economic losses^[3]^. ASF occurred as a rapid outbreak across the nation, indicating its highly contagious nature^[3]^. Given its similar clinical signs to several other porcine diseases^[1]^ and the lack of laboratory facilities for diagnosis on farms, diagnosing ASF often requires infected samples to be transported to a qualified laboratory, which delays diagnosis and increases the chances of contagion. Hence, accurate and on-site diagnosis of ASF is the key to controlling disease outbreaks.

Ours is the first report of a Cas12a-based assay using immunochromatographic lateral flow strips, termed CORDS, which enables the on-site detection of ASFV in one hour. This new assay requires no instrumentation other than a 37°C incubator, and its reaction temperature (body temperature) could be easily attained in an on-site situation. The whole reaction system comprises three tubes of lyophilized powder. Thus, the CORDS assay’s workflow only includes dissolution of lyophilized powders and addition of the samples, without any complex technical requirements. The whole detection process takes only one hour and displays its result through the band visualized on a lateral flow strip. Therefore, the CORDS assay has the advantage of detecting ASFV on-site (i.e., on farms or in slaughter houses) in a short time and without access to a diagnostics laboratory.

As a highly specific lateral flow strip-based assay, the specificity of CORDS is guaranteed by LbCas12a-crRNA targeting. LbCas12-crRNA can discriminate single nucleotide mutations such as point mutations in the seed region^[14, 15]^. In the present study, the CORDS assay did not cross-react with 13 nucleic acid samples from typical porcine viruses, making it superior to the methods reported until now^[7, 8]^. Thus, CORDS can distinguish ASF from other porcine diseases sharing similar clinical signs easily on site, even avoiding confusion between CSF and PRRS.

The LOD of the CORDS assay is at the femtomolar level for ASFV-specific NAs. To be precise, the system can detect ASFV p72 gene copies as low as 6×10^5^ copies/mL, which is more sensitive than isothermal amplification-based assays^[2]^ (which are considered good choices for on-site viral detection) and compares well with the newly reported lateral flow strip based assays for ASFV detection^[7, 8]^. The sensitivity of this assay is sufficient for ASF diagnosis on farms or in slaughter houses. During the ASFV infection process, blood virus levels can increase to 1 × 10^6^ copies/mL in 5–8 days post-infection (dpi)^[18]^. ASFV infections can result in virus levels of up to 1 × 10^9^ TCID_50_/mL in the blood^[4]^, indicating that virus genome copies can achieve > 1 × 10^9^ copies/mL. Hence, the femtomolar sensitivity of the CORDS assay should diagnose ASF at an early stage of infection (5–8 dpi) when the virus genome copy number reaches 1 × 10^6^.

For ease of use, the CORDS assay components are lyophilized in three tubes: the RAA mix, RAA buffer mix, and the Cas mix. We found that the assay maintained femtomolar sensitivity after lyophilization. Lyophilization significantly simplifies the operational process and improves the detection stability. According to our accelerated stability tests, the RAA buffer mix still maintained femtomolar sensitivity after storage at 37°C for one week. The lyophilized Cas mix should work well after storage at 4°C for one week without any sensitivity decrease. Therefore, these stability tests show that CORDS can be stored and transported under the conditions provided by the diagnostic kit’s manufacturers. It is envisaged that lyophilized CORDS has potential for industrial manufacturing and wide-spread on-site application. According to the accelerated stability tests, the lyophilized Cas mix is not as stable as the lyophilized RAA buffer mix and manufacturer-provided RAA mix. Considering Cas12a is a rather stable protein, the cause might be instability of the crRNA. RNA modification or capping may help to solve this problem and enable transportation at room temperature to lower the assay costs.

In addition to the lateral flow-strip readout, the RAA-Cas12a-based system also provides a fluorescent readout assay with high sensitivity. Our Cas12a-based nucleic acid fluorescence reporting system reached a sensitivity level of 1 × 10^-9^ M without amplification of dsDNA targets. In combination with RAA amplification, the RAA-Cas12a-fluorescence assay detected the dsDNA target at a sensitivity level of 10 aM, thereby reaching the level of other Cas12/13-based assays^[11-15]^. The fluorescent readout format also enables kinetic tests on reporter cleavage, providing information for reaction optimization and the possibility of more accurate detection. Compared with the well-establish real-time PCR, the RAA-Cas12a-fluoresence system is capable of high-throughput detection with lower costs and simpler operation.

To sum up, RAA-Cas12a-based system, especially the CORDS assay, is a promising ASFV detection method with high sensitivity and specificity, as well as having no need for scientific instruments and skilled operators. The lyophilized CORDS assay is now ready for industry manufacturing and wide-ranging applications.

## Acknowledgment

This work was supported by the National Natural Science Foundation of China (Grant 81202585) and Fundamental Research Funds for the Central Universities (2019MS089) from South China University of Technology.

